# Gut microbial dysbiosis in individuals with Sjögren’s disease

**DOI:** 10.1101/645739

**Authors:** Roberto Mendez, Arjun Watane, Monika Farhangi, Kara M. Cavuoto, Tom Leith, Shrish Budree, Anat Galor, Santanu Banerjee

## Abstract

**Purpose:** To evaluate the gut microbiome in individuals with Sjögrens and correlate bacterial profiles to dry eye (DE) measures.

**Methods:** Prospective case series of individuals with confirmed (n=13) and unconfirmed (n=8) Sjögrens (n=21; total cases) as compared to healthy controls (n=10). Stool was analyzed by 16S pyrosequencing and associations between bacterial classes and DE symptoms and signs were examined.

**Results:** Firmicutes was the dominant phylum in the gut, comprising 40-60% of all phyla. On a phyla level, subjects with Sjögrens (confirmed and unconfirmed) had depletion of Firmicutes (1.1- fold) and an expansion of Proteobacteria (3.0-fold), Actinobacteria (1.7-fold), and Bacteroidetes (1.3-fold) compared to controls. Shannon’s diversity index showed no differences between groups with respect to the numbers of different operational taxonomic units (OTUs) encountered (diversity) and the instances these unique OTUs were sampled (evenness). On the other hand, Faith’s phylogenetic diversity showed increased diversity in cases vs controls, which reached significance when comparing confirmed Sjögrens and controls (13.57 ± 0.89 and 10.96 ± 0.76, p=0.02). Using Principle Co-ordinate Analysis, qualitative differences in microbial composition were noted with differential clustering of cases and controls. Dimensionality reduction and clustering of complex microbial data further showed differences between the three groups, with regard to microbial composition, association and clustering. Finally, differences in certain classes of bacteria correlated with DE symptoms and signs.

**Conclusions:** Individuals with Sjögrens have gut microbiome alterations as compared to healthy controls. Certain classes of bacteria were associated with DE measures.

## Introduction

Sjögrens is a chronic autoimmune disease characterized by oral and ocular dryness. It is a common autoimmune disorder, affecting 0.5-4% of the population, with more than 2 million Americans living with the disease.^(1)^ Recently, there has been an interest in understanding interactions between gut bacteria and mucosal immunity in a number of eye diseases including Sjögrens.(2) In a homeostatic state, commensal bacteria serve as a metabolically active organ and aid the host in a plethora of activities.(3) For example, many plant polysaccharides cannot be directly digested and are instead transformed by gut bacteria into short-chain fatty acids (SCFAs), like acetic acid and butyric acid. (4) Interestingly, some SCFAs enhance the death of effector T cells and promote proliferation of regulatory T cells in the intestine, and thus help suppress inflammation and the development of auto-immune disease.(5) On the other hand, abnormal alterations in the gut bacterial community (dysbiosis) can have negative effects on the host.(6)

Individuals with auto-immune diseases have been found to have gut microbiome alterations compared to healthy controls, including individuals with spondyloarthritis, rheumatoid arthritis, Behçets, and Sjögrens.(7–10) In Sjögrens, greater relative abundances of *Pseudobutyrivibrio, Escherichia/Shigella* and *Streptococcus* and reduced relative abundances of *Bacteroides, Parabacteroides, Faecalibacterium* and *Prevotella* were noted compared to controls. Reduced gut microbiome diversity was also found to correlate with overall disease severity.(10) The association between gut bacteria and auto-immune disease is likely a two-way street. On one hand, gut microbiome abnormalities can lead to systemic inflammation and, conversely, systemic inflammation can preferentially deplete beneficial gut bacteria and promote the growth of commensal bacteria with potential pathogenic properties.(6, 11–14)

As there is limited data on gut microbial composition in Sjögrens, we performed this study to evaluate the diversity, dimensionality and constituency of the gut microbiome in individuals with Sjögrens in a South Florida population and to correlate gut microbiome profiles to clinical parameters of disease. Understanding the interactions between intestinal biodiversity and the immune system will be fundamental in deciphering and treating the pathogenesis and causes of autoimmune diseases, including eye diseases.(15)

## Methods

### Study population

Individuals seen between November 2017 and February 2018 at the Miami Veterans Affairs (VA) Hospital or Bascom Palmer Eye Institute with confirmed or unconfirmed Sjögrens were invited to participate. Individuals were considered as having confirmed Sjögrens if they fulfilled the American College of Rheumatology definition having 2 of 3 of the following: 1) positive serum anti-SSA (Ro) and/or anti-SSB (La) or positive rheumatoid factor (RF) and antinuclear antibody (ANA) ≥ 1:320; 2) ocular staining score ≥3; and 3) presence of focal lymphocytic sialadenitis with focus score ≥1 focus/4mm2 in labial salivary gland biopsies.(16) Individuals were considered as having unconfirmed Sjögrens if they had dry eye (DE) symptoms and one or more of the following: (1) ≥1 early Sjögrens marker positivity; (2) aqueous tear deficiency (ATD) defined as Schirmer score with anesthesia ≤5 in either eye; or (3) an auto-immune disease (e.g. rheumatoid arthritis).

### Ethical approval

The Miami VA and University of Miami Institutional Review Boards (IRB) approved the prospective evaluation of patients. Informed consent was obtained from all subjects and the study was adherent with the principles of the Declaration of Helsinki.

### Clinical metrics

Demographic information for each participant was collected including age, gender, race, ethnicity, past ocular and medical history and current medications.

### DE symptoms

Participants completed two standardized DE symptoms questionnaires: the Dry Eye Questionnaire 5 (DEQ5)(17) (score 0-22) and the Ocular Surface Disease Index (OSDI)(18) (score 0-100).

### DE signs

Participants underwent a complete ocular surface exam of both eyes in the following order:

1. Ocular surface inflammation via matrix metalloproteinase (MMP) 9 levels (Inflammadry, Quantel, San Diego, CA)(19) graded based on the intensity of the pink line (0 = no line, 1 = faint pink line, 2 = pink line, 3 = intense pink line).
2. Tear breakup time (TBUT) using fluorescein stain measured three times in each eye and averaged.
3. Corneal staining using fluorescein stain evaluated using the National Eye Institute (NEI) scoring scale which assesses 5 areas of the cornea on a 0-3 scale with a total score generated by summing the 5 section scores.(20)
4. Basal tear production after anesthesia placement (measured in mm at 5 minutes) using Schirmer’s strips.
5. Meibum quality evaluated after expression with intermediate pressure applied to the lower eyelid (0 = clear; 1 = cloudy; 2 = granular; 3 = toothpaste; 4 = no meibum extracted).

### Stool collection and analysis

All subjects were given a stool collection kit for at home collection. Stool samples were collected and placed in a glycerol suspension, homogenized, and sent to OpenBiome (Cambridge, MA). Specimens were then frozen at −80°C until analysis. Total DNA was isolated using Power-soil/fecal DNA isolation kit (Mo-Bio, Germantown, MD) as per manufacturer’s specifications. All samples were quantified using the Qubit^®^ Quant-iT dsDNA Broad-Range Kit (Life Technologies, Grand Island, NY) to ensure that they met minimum concentration and mass of DNA.(21) To enrich the sample for the bacterial 16S V4 rDNA region, DNA was amplified using fusion primers designed against the surrounding conserved regions that are tailed with sequences to incorporate flow cell adapters and indexing barcodes (Illumina, San Diego, CA). Each sample was PCR amplified with two differently barcoded V4-V5 fusion primers and were advanced for pooling and sequencing. For each sample, amplified products were concentrated using a solid-phase reversible immobilization method for the purification of PCR products and quantified by electrophoresis using an (Agilent, Santa Clara, CA) 2100 Bioanalyzer. The pooled 16S V4V5-enriched, amplified, barcoded samples were loaded into the MiSeq cartridge (Illumina Inc, San Diego, CA), and then onto the instrument along with the flow cell. After cluster formation on the MiSeq Instrument (Illumina, San Diego, CA), the amplicons were sequenced for 250 cycles with custom primers designed for paired-end sequencing.

In addition to patient samples, reagent controls were supplied in triplicate as background. Samples producing amplicons at later cycles compared to majority of samples were concentrated using Agencourt AMPureXP beads (Beckman Coulter, Indianapolis, IN). All samples were sequenced together after barcode-normalization subsequent to a preliminary sequencing run.

Using QIIME 2.0,(22) sequences were quality filtered and de-multiplexed using exact matches to the supplied DNA barcodes and primers. Resulting sequences were then searched against the SILVA database (v123) and clustered at 99% to obtain phylogenetic identities.

### Statistical analysis

OTU tables were rarefied to the sample containing the lowest number of sequences in each analysis. QIIME 2.0(22) was used to calculate alpha diversity and to summarize taxa. Descriptive statistics were used to describe relative compositions of bacteria on phyla, genera and class levels. Data were analyzed for significance by 2 tailed student t and Mann-Whitney U tests (GraphPad Prism 8).

### Principal Coordinate Analysis

Principal Coordinate Analysis (PCA) was done using observation ID level. The PERMANOVA test was utilized for finding significant whole microbiome differences among discrete categorical or continuous variables with randomization/Monte Carlo permutation test (with Bonferroni correction). The fraction of permutations with greater distinction among categories (larger cross-category differences) than that observed with the non-permuted data was reported as the p-value. False discovery rate (FDR) corrected p-value (q-value) <0.05 was considered significant across groups.

### Comparison of diversity indices

α-diversity matrices were compared between groups using Kruskal-Wallis pairwise rank tests or its variant, the Mann-Whitney U test.

### Dimensionality reduction and bacterial association analysis

We utilized both sequence-based and OTU-matrix dimensionality reduction and clustering algorithms. Compared to Qiime-derived PCA, which is done sample-wise, these algorithms are identity agnostic and decipher qualitative association (and disassociations) between experimental groups. The following methods were used:

a. **t-SNE** (t-Distributed Stochastic Embedding): t-SNE algorithm was implemented using group-wise OTU matrix with SeqGeq 1.5.0 software (FloJo LLC, Ashland, OR). t-SNE plots were generated with a perplexity value of 30 and 1000 iterations.
b. **Reference-independent binning**: This algorithm performs a reference-independent deconvolution of metagenomic sequences to reduce non-linear dimensionality of the samples. We used the java implementation of VizBin(23) for this analysis. We used a minimum contig length of 200 bases, k-mer length of 5, perplexity 30 with 1000 iterations for this study. Both individual samples and concatenated groups were analyzed.
c. **Bacterial association and clustering**: We used Graphia software (Edinburgh, UK) and its implementation of Markov Clustering algorithm (MCL).(24) Using our group-wise OTU matrix, MCL looked for cluster structures using mathematical bootstrapping. While this method is still agnostic to phylogenetic hierarchy in the matrix, we used the hierarchy as identifying markers to understand the bacterial clusters and changes in those clusters within the 3 study groups. MCL used the stochastic flow of the matrix to decipher the distances between the OTUs at equilibrium, thereby generating a cluster map by using correlation scores as distance. For generating the cluster, nodes scoring above a Pearson correlation value of 0.85 were used.

### Correlations between bacterial classes and clinical measures

Non-parametric Spearman rho (95% confidence interval) was calculated for each DE parameter and α<0.05 was considered to be significant in a 2-tailed test. Multivariable linear regression analyses were performed to assess these relationships while considering demographics.

## Results

### Study population

21 subjects were enrolled in the study (**Table 1**), 13 with confirmed Sjögrens and 8 with unconfirmed Sjögrens based on early marker positivity (n=2), aqueous tear deficiency (n=3), or presence of a systemic auto-immune condition (n=3). The mean age of the population was 60 years (range 33-71, standard deviation (SD) 8.8), 14 (67%) were female, 12 (57%) were white, and 8 (38%) were Hispanic. Co-morbidities included diabetes (n=2), hypothyroid (n=5), hypertension (n=8), sleep apnea (n=4), rheumatoid arthritis (n=5), psoriatic arthritis (n=1), systemic sclerosis (n=1), and systemic lupus erythematosus (n=1). The majority of individuals reported DE symptoms (mean DEQ5 11.6, mean OSDI 41) and had corneal staining (mean 7.2) (Table 1). Controls consisted of 10 individual samples provided by OpenBiome. The mean age of the controls was 26 (SD=5.6) with all controls being male. Cases with Sjögren’s (n=21) were older than controls (59 vs 26, p=0.07).

**Table 1:** Clinical characteristics of the study population

| Variable | No. of Patients | % |
| --- | --- | --- |
| **Demographics** |  |  |
| Age, years* | 58.7 ± 8.8 (33-71) |  |
| Gender, male | 7 | 33 |
| Race, white | 12 | 57 |
| Ethnicity, Hispanic | 8 | 38 |
| Smoking |  |  |
| Past | 3 | 14 |
| Current | 1 | 5 |
| **Dry eye symptoms** |  |  |
| DEQ5 | 11.6 ± 4.8 (0-18) |  |
| OSDI | 41.2 ± 22.6 (0-83.3) |  |
| **Dry eye signs<sup>†</sup>** |  |  |
| Inflammation via Inflammadry | 1.3 ± 1.1 (0-3) |  |
| Tear break up time | 5.7 ± 2.9 (2-13) |  |
| Corneal staining | 7.2 ± 3.6 (1-14) |  |
| Schirmer score | 8.1 ± 6.1 (1-25) |  |
| Meibum quality | 2.4 ± 0.9 (0-4) |  |

DEQ5=Dry Eye Questionnaire 5; OSDI=Ocular Surface Disease Index
\*mean ± standard deviation (range)
†more abnormal value between the two eyes

### Gut microbial landscape in Sjögrens compared to controls

Firmicutes was the dominant phylum in the gut in all individuals, composing between 40-60% of all phyla, followed by Bacteroidetes. Cases (confirmed + unconfirmed) had a depletion of Firmicutes (1.1- fold), and an expansion of Proteobacteria (3.0-fold), Actinobacteria (1.7-fold) and Bacteroidetes (1.3-fold), compared to controls **(Figure 1A)**. There was no perceivable difference between the cases and controls in terms of Bacteroides-Firmicutes ratio, although a lower ratio is considered a hallmark of inflammatory disease (**Figure 1B**). While Clostridia, Bacteroidea and Actinobacteria were the dominant classes in all groups (**Figure 1C**), Actinomycetaceae (3.6 fold, p=0.01), Eggerthellaceae (6.2 fold, p=0.001), Lactobacillaceae (8.8 fold, p = 0.02), Akkermanciaceae (4.7 fold, p=0.04), Coriobacteriaceae (2.5 fold, p=0.04) and Eubacteriaceae (7.4 fold, p=0.02) had significantly increased abundance in cases compared to controls.

**Figure 1:**
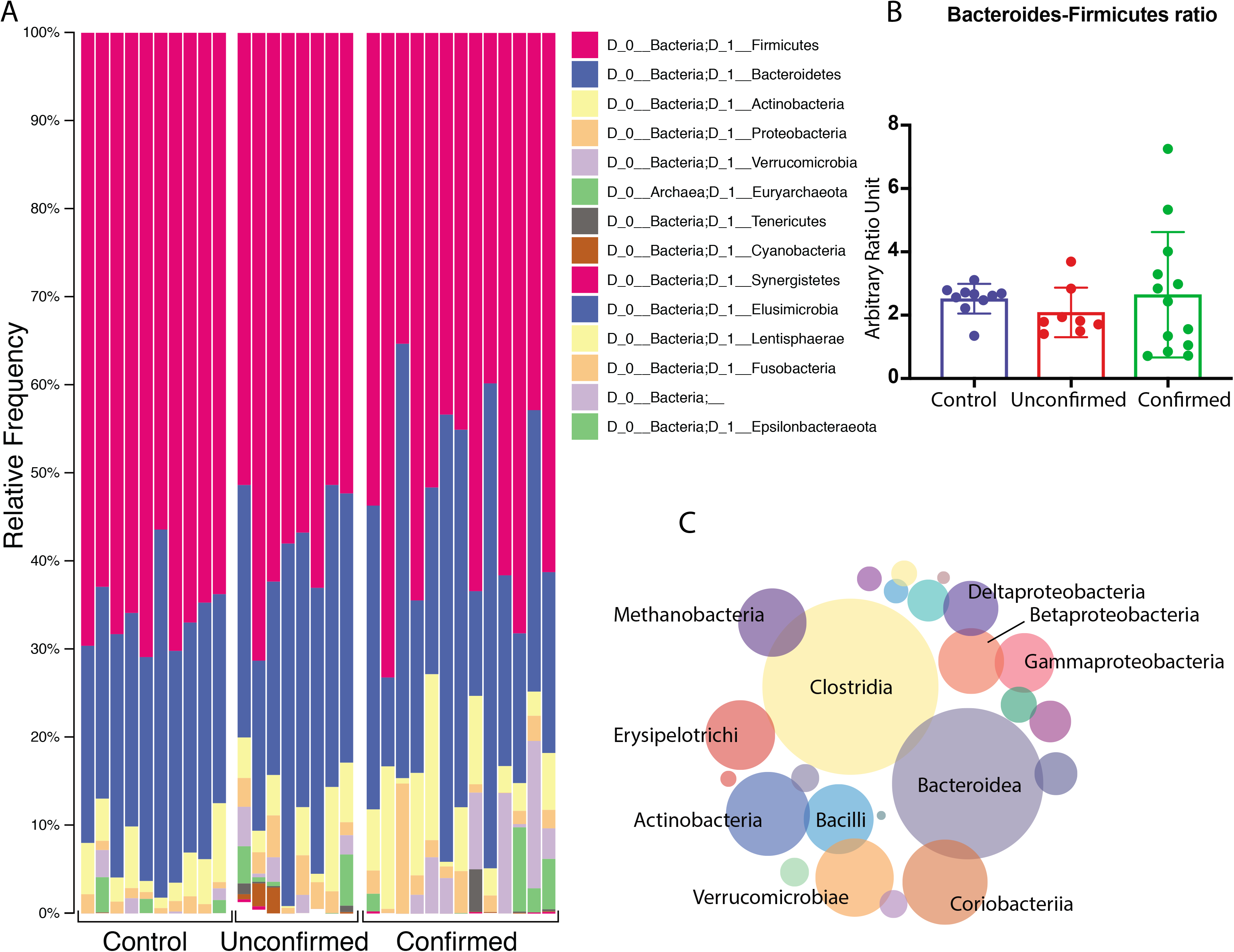
Overall distribution of bacterial Phyla and Classes in Control and Sjögrens (Unconfirmed and Confirmed) gut microbiome. (**A**) All three study groups exhibit a Firmicutes-Bacteroidetes dominated microbiome, with significant presence of Actinobacteria and Proteobacteria. (**B**) Bacteroidetes-Firmicutes ratio shows an upward trend for confirmed Sjögrens group, but it is statistically insignificant. (**C**) Dominant representative bacterial classes among all study subjects across the three study groups.

### Distance matrices show significant differences between controls, unconfirmed and confirmed Sjögrens gut microbiota

Bray-Curtis Principle Co-ordinate Analysis (pCoA; beta-diversity, **Figure 2A**) was used to qualitatively examine differences in microbial composition. There was a distinct clustering of the controls compared to cases (blue, circled). As mentioned before, there was a distinct age difference between controls and cases, with 2 individuals each from cases and controls overlapping in age (33-40 years, marked with arrows). From pCoA analysis, there was no evident similarity in their gut microbial composition. Next, unifrac distances were measured (**Figure 2B**) between the three groups and a pairwise PERMANOVA test was performed with false discovery rate correction (**Figure 2C**; FDR; q-value). Both Sjögrens groups exhibited significant compositional changes compared to controls, while these changes were insignificant when comparing the confirmed and unconfirmed cases. PERMANOVA test showed decrease in genera *Faecalibacterium* and *Viellonella*, classes Ruminococcaceae and Lachnospiraceae, and orders Clostridiales and Bacteroides comparing controls to unconfirmed to confirmed Sjögrens. There was also an increase in the genera *Megasphaera, Parabacteroides* and *Prevotella* (**Figure 2D**). These differences were major contributors to the significant differences seen in Figure 2A-C.

**Figure 2:**
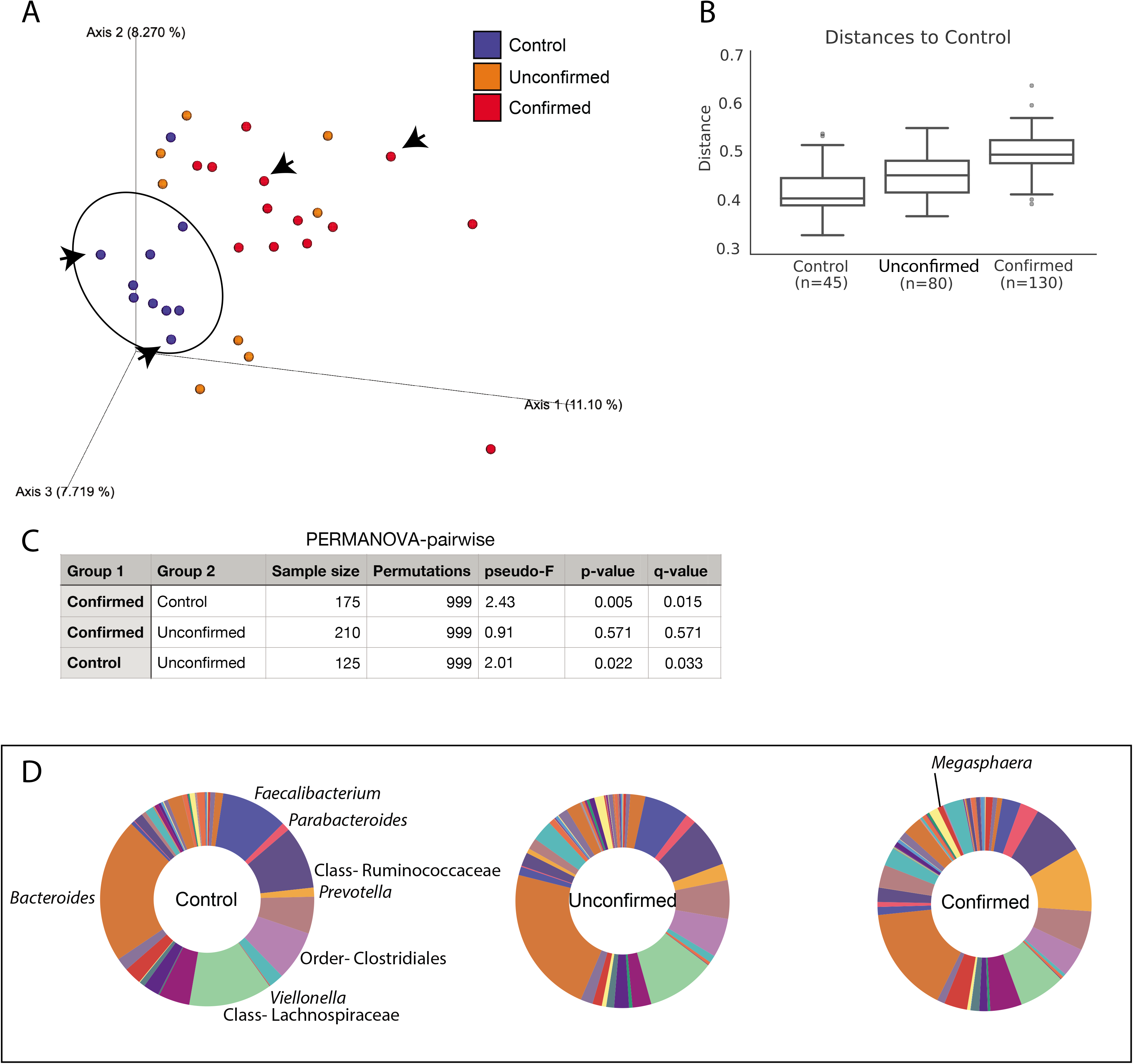
Microbial differences between Controls and Sjögrens. (**A**) Bray-Curtis Principal co-ordinate analysis (pCoA) showing the distribution of the three study groups. Controls (circled) cluster distinctly compared to Sjögrens cases. Arrows point to patients with overlapping age among Controls and Cases suggesting that microbial changes are not influenced by the age differences between patients. (**B**) Box and whisker plot showing all three study groups and (**C**) pairwise PERMANOVA on the UniFrac distances (unweighted) showing significant differences between Controls and each Sjögrens group (Unconfirmed and Confirmed). Compositional differences between Unconfirmed and Confirmed Sjögrens are not significant. (**D**) PERMANOVA test further shows the major microbial components within the three study groups driving the significance above. Genera are italicized and upper hierarchical groups are labeled.

### Comparison of diversity matrices between controls, unconfirmed and confirmed Sjögrens gut microbiota

Next, we looked at the different diversity matrices to understand the distribution of OTUs across all three groups. We started with the Shannon’s index, (25, 26) which accounts for both the abundance and evenness (measure of species diversity and distribution in a community) in all three groups (**Figure 3A**). Using Shannon’s index, we noted no differences between the overall diversity within groups, implying that the numbers of different OTUs encountered (diversity), as well as the instances these unique OTUs were sampled (evenness) were similar between control, unconfirmed and confirmed Sjögrens samples. Faith’s phylogenetic diversity (Faith’s PD) represents taxon richness of each sample, expressed as the numbers of phylogenetic tree branches encountered for each sample.(27) Faith’s PD data (**Figure 3B**) showed that microbial diversity was higher in cases compared to controls and the differences attained statistical significance between the confirmed Sjögrens and control groups. Unlike Shannon’s index, Faith’s PD is informed by OTU identities. This essentially means that while richness and evenness (Shannon’s) is similar among the groups, diverse bacterial groups replace existing ones, thereby enriching phylogenetic diversity in cases compared to controls. As we looked deeper into changes within cases, we did not find any significant differences, either in major diversity indices or within major phyla distribution between unconfirmed and confirmed Sjögrens (**Table 2**).

**Figure 3:**
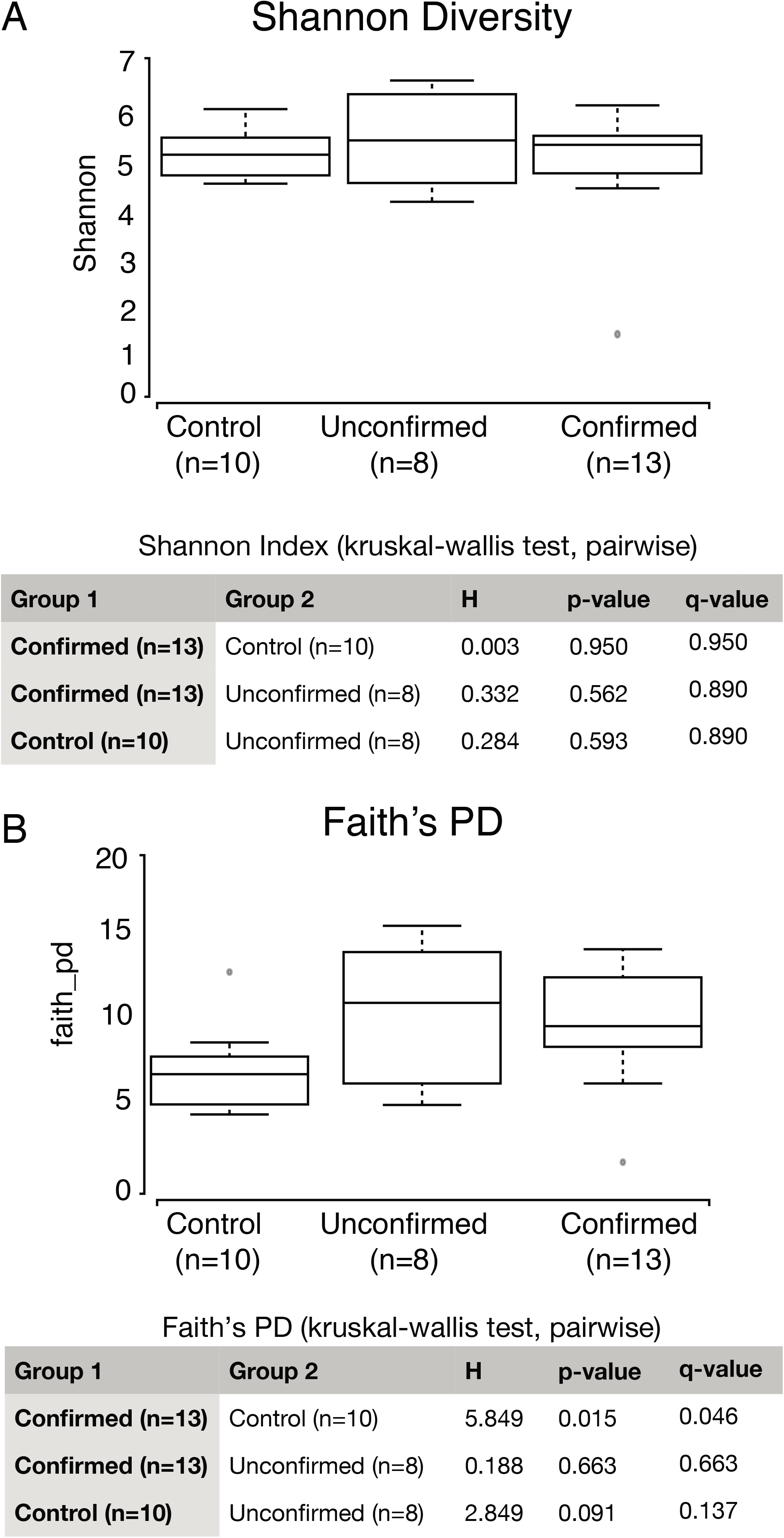
Microbial diversity between Controls and Sjögrens. Among other α-diversity matrices shown in Table 2, two of the major indices (**A**) Shannon and (**B**) Faith’s PD are displayed in the figure. While Shannon’s diversity did not show any differences between the groups, Faith’s PD index showed significant differences between Controls and Confirmed Sjögrens groups.

**Table 2:** Comparison between subjects with Confirmed vs Unconfirmed Sjögrens

| **Diversity indices/**<br>**Phylum/ratio** | **Confirmed**<br>**Mean (SD)** | **Unconfirmed**<br>**Mean (SD)** | **p-value** |
| --- | --- | --- | --- |
| Chao1 diversity | 116 (30) | 118 (35) | 0.88 |
| Faith's phylogenetic diversity | 57.5 (10.7) | 60.7 (11.2) | 0.52 |
| Shannon's index | 4.7 (0.64) | 4.9 (0.68) | 0.67 |
| Observed OTU | 891 (186) | 954 (155) | 0.44 |
| Simpson diversity | 0.076 (0.04) | 0.073 (0.04) | 0.90 |
| Actinobacter | 71 (70) | 45 (36) | 0.28 |
| Bacteroides | 332 (230) | 577 (744) | 0.39 |
| Firmicutes | 3502 (1898) | 4736 (1184) | 0.12 |
| Proteobacteria | 3.8 (10.0) | 1.3 (1.8) | 0.50 |
| Firmicutes/Bacteroides | 27 (58) | 30 (38) | 0.92 |

OTU=Operational taxonomy unit; SD=standard deviation

### Group-wise dimensionality reduction shows differential clustering between controls and both case groups

We implemented non-linear dimensionality reduction on the microbiome data using both group-wise OTU matrix (**Figure 4A**) and group-wise concatenated raw sequences (**Figure 4B**). As mentioned in the methods section, these plots are phylogeny and sample ID agnostic and define a similar probability distribution of all OTUs within a group.(28) Similar parameters were used to generate a two-dimensional density plot for controls, unconfirmed and confirmed Sjögrens samples. As seen in **Figure 4A**, control OTUs show a definite pattern of clustering that differs from the unconfirmed Sjögrens group. The differences encompass appearance of distinct clusters, dissociation of coalesced clusters compared to controls and disappearance/coalescence of distinct clusters compared to controls. Differences in these patterns were also seen in the distribution of OTUs in the confirmed Sjögrens group. Compared to controls, a distinct strengthening of existing clusters was seen accompanied by emergence of novel clusters, which is corroborative of increase in relative abundance of distinct classes/genera of bacteria in both case groups (**Figure 2D**). Differences in the population characteristics were even more evident when we did a reference-independent deconvolution of bacterial sequences as shown in **Figure 4B**.(23) In controls, there was an amorphous distribution of sequences with 10 distinct major clusters, several minor clusters and numerous un-clustered sequences in the middle. This pattern differed dramatically in unconfirmed Sjögrens group, in which there was a major rearrangement of the sequences into major clusters, with reduction in minor and un-clustered sequences. This pattern differed further in the confirmed Sjögrens group, where more similar sequences cluster into increased numbers of major clusters, at the expense of un-clustered sequences and minor clusters.

**Figure 4:**
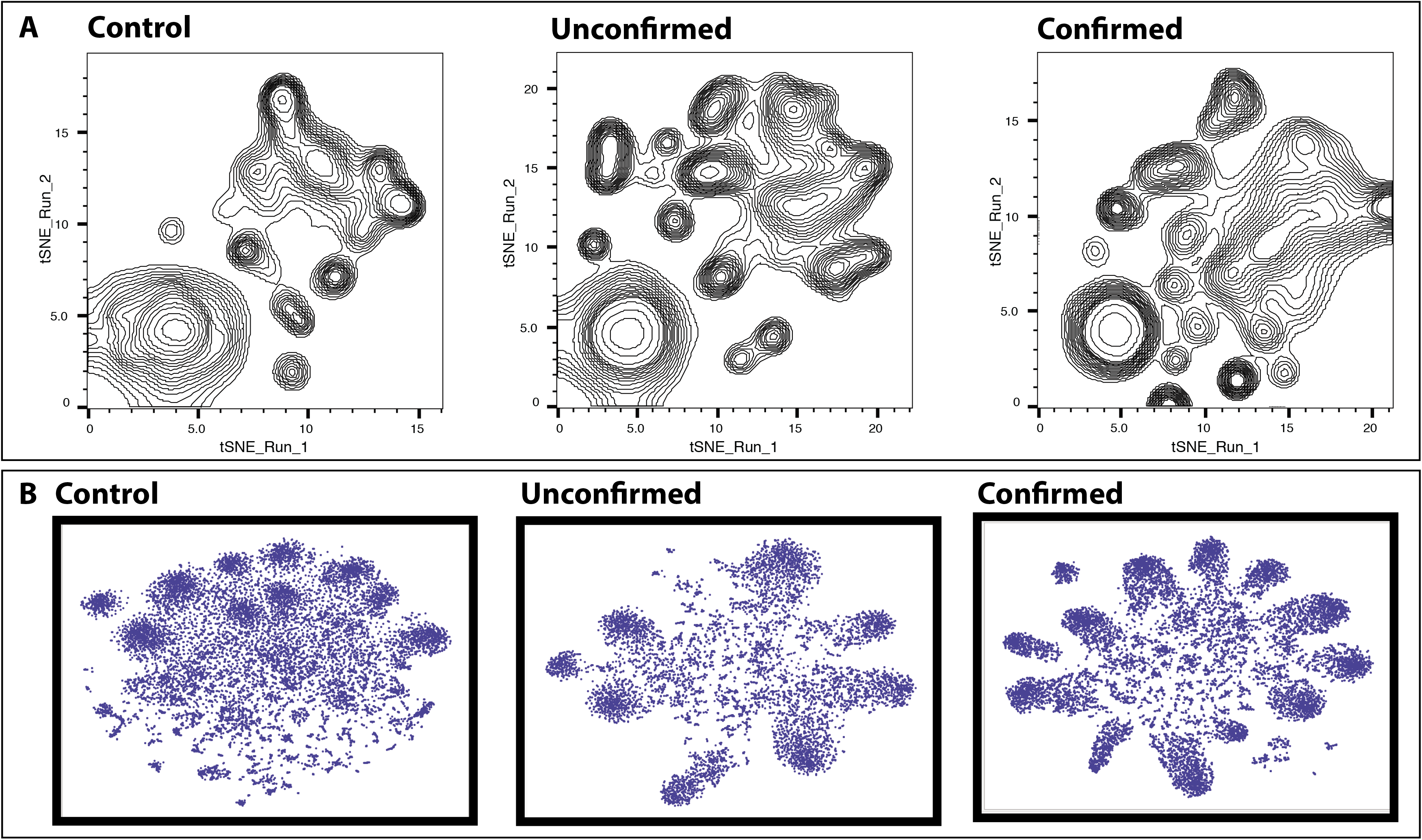
Dimensionality reduction of microbiome data and differential clustering within the three study groups. (**A**) t-Distributed Stochastic Embedding (t-SNE) implemented on group-wise OTU matrix for the three study groups. Figure shows control OTUs show a definite pattern of clustering that differs from the Unconfirmed and Confirmed Sjögrens. (**B**) Reference-independent deconvolution of bacterial sequences divided by the three study groups. While there are distinct differences between Unconfirmed and Confirmed Sjögrens in terms of clustering and distribution of individual sequences, the differences are dramatic compared to Controls.

### Bacterial association networks significantly change in cases compared to controls

Bacterial association was calculated using Markov Clustering Algorithm (MCL), which calculates correlations of vector abundance of OTUs within each group. This results in clustering of OTUs that are most likely to co-occur together in a network, using correlation values as the distance matrix. For example, the higher the correlation value between 2 OTUs, the more likelihood that they would associate within a cluster. This also allows for each OTU to participate in multiple clusters. As shown in **Figure 5**, control OTUs exhibited a single large super-cluster composed of 3 major clusters and several minor independent clusters. In unconfirmed Sjögrens, major constraints have been introduced into the network structure with the emergence of more clusters within the major super-cluster. In confirmed Sjögrens, these constraints seemed to be exacerbated, as the super-cluster stretched and expanded and new independent clusters emerged. A comparison of the major clusters in control group to unconfirmed Sjögrens showed a major rearrangement of bacterial association within its major clusters (**Supplementary Figure 2 for controls, Supplementary Figure 3 for unconfirmed and Supplementary Figure 4 for the Confirmed group**). This included the increase in numbers of clusters, accompanied by addition of phyla representation in major clusters. Furthermore, within each phylum, it was evident that the genera representation differed compared to controls (e.g. expanded representation of the genus *Prevotella* in unconfirmed Sjögrens clusters and the phylum Actinobacteria having different genera in each of the three groups). As evident from **Figure 5**, confirmed Sjögrens clusters have increased in numbers with major clusters composed of more diverse phyla with disparate genera representation.

**Figure 5:**
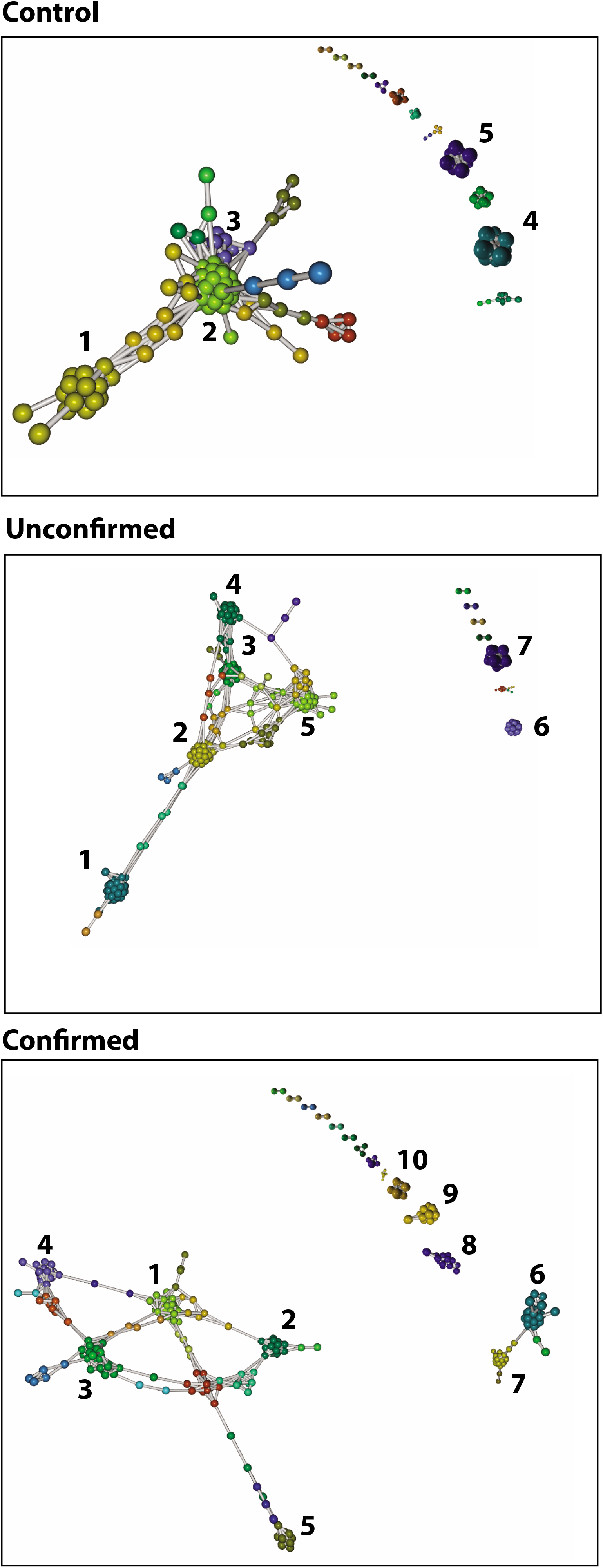
Markov Clustering algorithm and bacterial associations within the three study groups. As shown, control OTUs exhibited a single large super-cluster composed of 3 major clusters and several minor independent clusters. In Unconfirmed Sjögrens, major constraints have been introduced into the network structure with the emergence of more clusters within the major super-cluster. In Confirmed Sjögrens, these constraints seemed to be exacerbated, as the super-cluster stretched and expanded and new independent clusters emerged. Identities of the microbes comprising each cluster within the three groups is given in Supplementary sheet 1.

### Dry eye parameters correlate with several bacterial classes

Since unconfirmed and confirmed Sjögrens patients exhibited altered gut microbiota compared to controls, we evaluated several DE parameters and their correlation to changing bacterial classes in the two case groups compared to controls. As shown in **Table 3**, several bacterial classes exhibited positive or negative correlation to symptoms (DEQ5, OSDI) and signs (ocular surface inflammation, corneal staining, tear production). This relationship was unchanged when considering the effects of age, gender, ethnicity and race in a multivariable model.

**Table 3:** Correlations between clinical signs and gut microbial classes, controlling for demographics.

| **Confirmed Sjögren's** | **Class** | **rho** | **P (two tailed)** | **Comparison with the literature** |
| --- | --- | --- | --- | --- |
| DEQ5 | Barnesiellaceae | 0.65 | 0.02 | ↑ in JIA (29) |
|  | Muribaculaceae | -0.57 | 0.04 |  |
|  | Ruminococcaceae | 0.70 | 0.009 | ↓ in IBD and psoriasis (30) |
| OSDI | Atopobiaceae | -0.65 | 0.02 |  |
|  | Rikenellaceae | 0.56 | 0.048 | ↑ in SLE mice (31) |
|  | uncultured | -0.58 | 0.04 |  |
| Ocular surface inflammation* | Christensenellaceae | 0.66 | 0.03 |  |
| Corneal staining* | Christensenellaceae | 0.65 | 0.02 |  |
|  | Clostridia Family XIII | 0.63 | 0.04 | ↑ in RA and IBD-arthritis (48) |
|  | Enterobacteriaceae | -0.74 | 0.008 | ↑ in MS (49) |
| Schirmer's score* | uncultured | -0.67 | 0.04 |  |
|  | Victivallaceae | 0.67 | 0.03 | ↓ in psoriasis (50) |
| **Unconfirmed Sjögren's** | **Class** | **rho** | **P (two-tailed)** |  |
| DEQ5 | Clostridia Family XI | -0.87 | 0.009 | ↑ in RA and IBD-arthritis (48) |
| OSDI | Coriobacteriaceae | 0.90 | 0.005 | ↑ in SLE mice (51) |
|  | Defluviitaleaceae | 0.75 | 0.04 |  |
| Ocular surface inflammation* | Christensenellaceae | 0.80 | 0.04 |  |
|  | Ruminococcaceae | 0.90 | 0.02 | ↓ in IBD and psoriasis (30) |
| Corneal staining* | Clostridia Family XIII | 0.87 | 0.02 | ↑ in RA and IBD-arthritis (48) |
|  | Bacteroidaceae | 0.83 | 0.03 | ↓ in MS (49) |
|  | Pasteurellaceae | 0.93 | 0.007 |  |

DEQ5=Dry Eye Questionnaire 5; OSDI=Ocular Surface Disease Index; JIA= Juvenile Idiopathic Arthritis; RA=Rheumatoid arthritis; IBD=Inflammatory bowel disease; SLE=Systemic Lupus Erythro; MS=Multiple Sclerosis
\*For all dry eye signs, value from more severely affected eye used in the analysis

## Discussion

We demonstrated that individuals with Sjögrens had gut microbiome alterations compared to controls. In our study, we found that cases had more diverse phyla with disparate genera representation compared to controls. Specifically, these changes included a decrease in genera *Faecalibacterium* and *Viellonella*, classes Ruminococcaceae and Lachnospiraceae, and orders Clostridiales and Bacteroides from controls to unconfirmed, and then from unconfirmed to confirmed Sjögrens and an increase in the genera *Megasphaera*, *Parabacteroides* and *Prevotella*. Similar changes in microbial association and networks have been shown to alter virulence and metabolic behavior of microbes in other disease models(29–31) and hence, assumes added importance in the role of host-microbiota interactions in health and disease. In addition, we expanded our exploration of composition beyond diversity to evaluate clustering patterns in disease, noting differences between controls, unconfirmed and confirmed Sjögrens with respect to pattern and clustering due to relative perturbations in bacterial populations. These changes in community behavior align with biological context, which demonstrate that compositional changes in microbiota due to disease is not a singular change. Changes in relative abundance of microbial species results in imposition of constraints on co-occurrence network(s), resulting in induction/expulsion of other species, or complete rearrangement of networks.(6)

Our findings demonstrate both similarities and differences compared to prior data in Sjögrens. In a study of 10 individuals with Sjögrens, greater relative abundances of *Pseudobutyrivibrio, Escherichia/Shigella, Blautia* and *Streptococcus* and lower abundances of *Bacteroides, Parabacteroides, Faecalibacterium* and *Prevotella* were noted compared to 45 healthy controls identified from the Human Microbiome Project.(10) Similar to our population, controls were significantly younger than cases (27 ± 5 years old vs 59 ± 14 years old). A commonality in both studies was the presence of dysbiosis in Sjögrens associated DE compared to controls, albeit with differences in bacterial signatures. Both studies noted a decrease in relative abundance of *Faecalibacterium* and *Bacteroides* in Sjögrens, but in our study, we saw an increase in relative abundance of *Prevotella*, a bacteria implicated in rheumatoid arthritis.(32, 33) Another difference was in phylogenetic diversity, in which we found increased diversity in individuals with Sjögrens compared to controls whereas the former study found a significant inverse correlation between diversity and disease severity (r = - 0.72, P = 0.01). Several differences must be considered when interpreting results between the two studies including differences in hypervariable region targets (V1-3 vs V4-5), curation status (2016 vs 2018) confidence interval (CI) of OTU database (97% vs 99%), controls (Human Microbiome Projects vs OpenBiome Stool Bank), geographical location (Texas vs Florida), and indices used (absolute OTU counts vs Faith’s PD). As a demonstration of how these differences can effect results, when we reanalyzed our data using the SILVA (v123) 97% CI database, the slight but significant increase in Faith’s PD in confirmed Sjögrens vs controls was lost (**Supplementary Figure 1**). As Faith’s PD is a measure of number of nodes in a phylogenetic tree, it is understandable that values would change with a lower number of hits.

Our findings mirror changes noted in other autoimmune conditions, including ones related to Sjögrens. Similar to our data, increased abundance of Actinobacteria,(34) specifically *Eggerthella* and *Actinomyces, Prevotella copri*,(33) *Lactobacillus*(35) and decreased abundance of *Faecalibacterium*,(34) *Bacteroides*,(8) *Lachnospiraceae*(33) and *Clostridiales*(33) were reported in individuals with rheumatoid arthritis as compared to controls. Interestingly, some of these dysbiotic signatures normalized with anti-inflammatory therapy.(36) In a similar manner, *Viellonella* was reduced in ankylosing spondylitis(37), *Ruminococcaceae* was reduced in inflammatory bowel disease and psoriasis(30) and *Megasphaera* was increased in primary biliary cirrhosis,(38) all mirroring our findings in Sjögrens. Beyond composition and not tested herein, bacterial metabolites of individuals with autoimmune disease has been found to differ from controls. For example, individuals with Behçets had less butyrate production in their gut compared to controls.(9) A similar findings was indirectly noted in Sjögrens with a 50% decrease in relative abundance of OTUs classified to the high butyrate producer *Faecalibacterium prausnitzii* compared to controls.(10)

These dysbiotic signatures may have a causal role in Sjögrens associated DE. Inflammation is a hallmark of DE in individuals with and without Sjögrens(39, 40) and it is well-established that the gut microbiome has an impact on inflammation and immunity.(6, 11–14) The commensal gut microbiome monitors mucosal immunity through the generation of anti-inflammatory regulatory T cells (Treg cells) and pro-inflammatory Th17 cells. The balance between these cells protects the mucosa from pathogenic microorganisms and limits excessive T cell responses via key mediators, including TGF-B, IL-6, retinoic acid and SCFA. For example, specific *Clostridia* species have been found to specifically induce Th17 cells in the small intestine and in extraintestinal sites during autoimmune inflammation. Other *Clostridia* clusters have been shown to induce Tregs and produce SCFAs to support Treg development.(41) In a similar manner, *Bacteroides* species can express polysaccharide A which suppresses Th17 inflammatory responses, allowing mucosal tolerance and subsequently colonization.(41) Putting this in context of our findings, reductions in commensals such as Clostridiales and *Bacteroides* may have an impact on the balance of Th17 and Treg cells, tipping the body towards auto-immunity.

Specific to the eye, altering the intestinal microbiome has been shown to influence eye disease. For example, CD25 knock out (KO) mice spontaneously develop DE and thus serve as a model of Sjögrens associated DE. Germ-free CD25KO mice had a worse DE phenotype compared to CD25KO control mice, including increased lacrimal gland inflammation and IFN□-producing T cells. Interestingly, recolonization of the gut microbiome improved the DE phenotype, with decreased lacrimal gland inflammation, IFN-□producing T cells and corneal staining.(42) Similar findings have been seen in other mice models. Desiccating stress applied to the ocular surface with a fan in germ-free mice led to corneal staining and ocular surface inflammation, resulting in a worse DE phenotype compared to conventionally house mice.(43) In another experiment, antibiotics administered in addition to desiccating stress reduced Bacteroidetes and Firmicutes and increased Proteobacteria in the gut and concomitantly caused a more severe DE phenotype compared to desiccating stress alone.(44) These experiments reinforce the idea of a gut-eye axis and highlight the possibility of gut microbiome modulation and a therapeutic approach in DE.

Our findings should be interpreted bearing in mind our study limitations, which included cases consisting of both confirmed and unconfirmed Sjögrens. The rationale for including both groups is that Sjögrens is often diagnosed late in the disease course as the traditional markers, SSA and SSB, become positive years after disease initiation, if at all.(45) As such, many individuals with DE that have a specific profile (aqueous tear deficiency, early marker positivity, DE in the setting of an established auto-immune disease such as rheumatoid arthritis) are considered as having Sjögrens-like DE but do not fit the ACR criteria for disease. In this study, we were interested in understanding gut microbiome profiles in both groups, although we acknowledge that the unconfirmed group likely has a more heterogeneous makeup. Fortunately, as evident from our PCA plot, data from our diverse patient population is driven into a tight cluster, suggesting significant disease-mediated microbial changes in both groups, compared to controls. In addition, we chose controls provided by a stool bank (Openbiome) so as to compare our population to a well-phenotyped, healthy control group, which differed significantly in age and varied in gender. An issue with contemporary controls (e.g. age matched veterans) is that other co-morbidities may affect microbiome health. As such, we modeled our work on prior studies, in which a similar approach also resulted in an age differential between cases to controls.(10) While age related differences in the gut microbiome have been noted when comparing very young children to adults, it seems that microbiome stabilizes to an adult-like composition by age 5.(46, 47) In addition, 16s rRNA sequencing method has limited genera coverage, which limits a detailed study of the microbiome. Finally, our study did not measure metabolic products of bacteria, such as butyrate, which would be indicative of the function of the microbiome.

Despite these limitations, our study findings are important as they set the foundation for modulating the gut microbiome as a potential therapeutic approach in Sjögrens associated DE. There are several ways to modulate the gut microbiome, including dietary intake, probiotics and fecal microbial transplantation (FMT). For example, FMT was used to modulate the gut microbiome in Graft Versus Host Disease (GVHD), another condition associated with DE. In four individuals with GVHD, FMT increased abundances of the beneficial bacteria *Lactobacillus, Bacteroides, Bifidobacterium* and *Faecalibacterium*, and concomitantly improved gastrointestinal symptoms such as defecation consistency and frequency. Future studies are needed to translate these findings to Sjögrens associated DE.

---

**Supplementary Figure 1:** Faith’s PD measurement between Controls, Unconfirmed Sjögrens and Confirmed Sjögrens when clustered against 97% CI and 99% CI SILVA database. (**A**) Dataset clustered against 97% CI-- Faith’s PD index is similar between the groups. (**B**) Dataset clustered against 99%CI— confirmed Sjögrens samples show incremental, yet significant enhancement in phylogenetic index, compared to controls. The test of significance used was a pairwise Mann-Whitney U test.

**Supplementary Figure 2:** Spreadsheet with the control group clusters with bacterial identities comprising each cluster marked in Figure 5.

**Supplementary Figure 3:** Spreadsheet with the Unconfirmed group clusters with bacterial identities comprising each cluster marked in Figure 5. As evident from this list, there are changes in phyla representation in major clusters.

**Supplementary Figure 4:** Spreadsheet with the Confirmed group clusters with bacterial identities comprising each cluster marked in Figure 5. Within each phylum genera representation differed compared to controls (e.g. expanded representation of the genus *Prevotella* in unconfirmed Sjögrens clusters and the phylum Actinobacteria having different genera in each of the three groups). As seen in Figure 5, confirmed Sjögrens clusters have increased in number with major clusters composed of more diverse phyla with disparate genera representation.

## Notes

#### Summary of Updates

Changes to supplementary figure

