## Supplementary figures and images for "Gut microbial dysbiosis in individuals with Sjögren’s disease"

### Supplementary Figure 1

#### Supplementary Figure 1

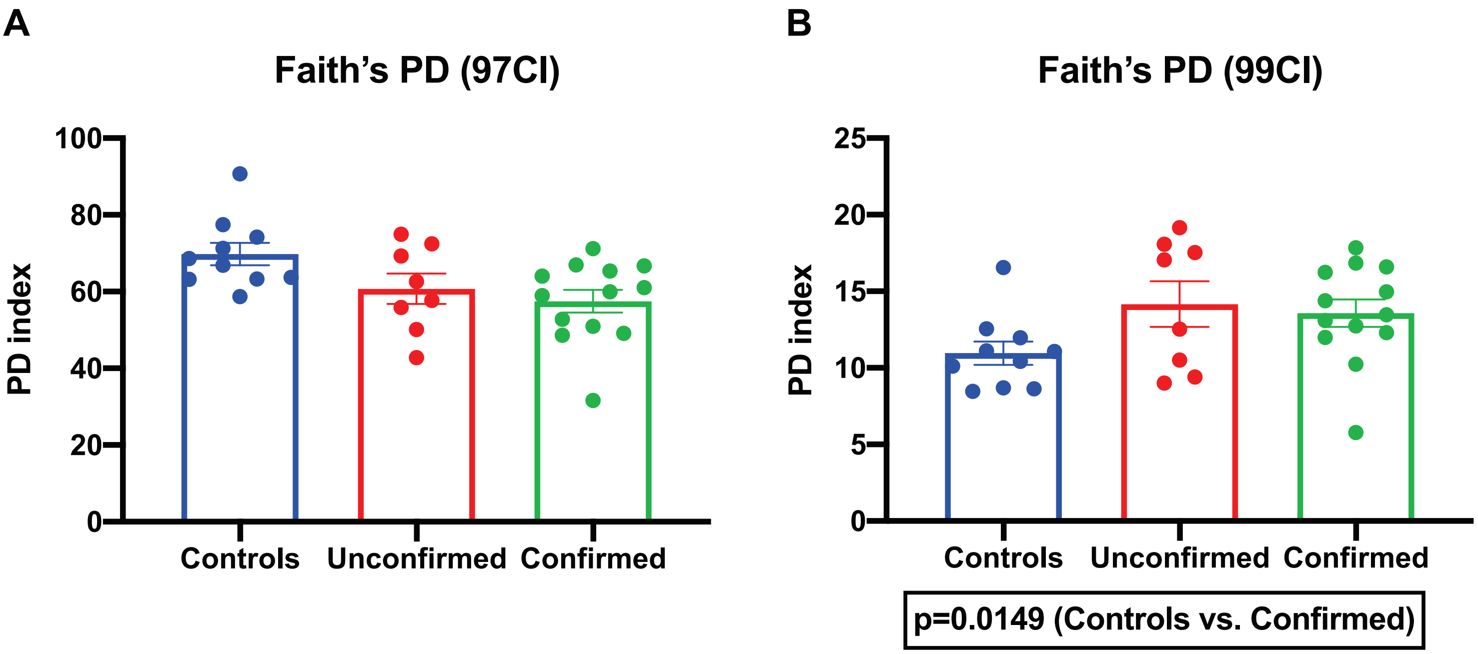
